# Mutualistic network architecture and eco-evolutionary feedbacks modulate the occurrence of transitions and stability in response to rising temperature

**DOI:** 10.1101/2023.10.08.561385

**Authors:** Gaurav Baruah, Tim Lakämper

**Affiliations:** Faculty of Biology, Theoretical Biology, University of Bielefeld, 33501 Bielefeld, Germany

**Keywords:** Individual variation, nestedness, connectance, mutualistic networks, eco-evolutionary dynamics, critical transitions

## Abstract

1. Ecological networks comprising of mutualistic interactions can suddenly transition to undesirable states, such as collapse, due to small changes in environmental conditions such as a rise in local environmental temperature.

2. However, little is known about the capacity of such interaction networks to adapt to changing temperatures and thereby impact the occurrence of critical transitions.

3. Here, combining quantitative genetics and mutualistic dynamics in an eco-evolutionary framework, we evaluate the resilience of mutualistic networks to critical transitions as environmental temperature increases. Specifically, we model the dynamics of a phenological optimum trait that determines the tolerance to local environmental temperature as well as temperature-dependent species interaction and evaluate the impact of trait variation and evolutionary dynamics in the occurrence of tipping points and community collapses.

4. We found that mutualistic network architecture, i.e., community size and the arrangement of species interactions, interacted with evolutionary dynamics to impact the onset of network collapses. In addition, some networks had more capacity to track the rise in temperatures than others and thereby delay the occurrence of threshold temperatures at which the networks collapsed.

5. However, such a result was modulated by the amount of heritable trait variation species exhibited, with high trait variation in the mean optimum trait value delaying the environmental temperature at which the network collapses.

6. Our study argues that mutualistic network architecture modulates the capacity of networks to adapt to changes in temperature and thereby impact the occurrence of community collapses.

## 1 Introduction

Mutualistic networks may suddenly transition from one stable state to an alternative undesirable state in response to changes in the external environment (Lever *et al*. 2014, Bascompte & Scheffer 2023, Baruah 2022, Jiang *et al*. 2018). Such networks comprising of positive interactions play a crucial role in maintaining biodiversity and providing beneficial ecosystem functions and services (Bascompte & Jordano 2007, Encinas-Viso *et al*. 2012, Bastolla *et al*. 2009, Schleuning *et al*. 2015). These networks consists of groups of species that rely on mutually positive interactions such as plant-pollinator interactions, seed dispersal etc. These mutualistic networks could undergo changes in strength of species interaction in response to changes in the external environment, such as rise in temperature or habitat destruction (Bhandary *et al*. 2023, Bascompte & Stouffer 2009, Revilla *et al*. 2015, Prakash & de Roos 2004, Baruah 2023, De Laender *et al*.). Due to rise in temperature, mutualistic interactions could be disrupted, as mismatches in plant-pollinator phenology for instance could occur, leading to the weakening of interactions (Hegland *et al*. 2009, Ibáñez *et al*. 2010, Gordo & Sanz 2005, Memmott *et al*. 2007). Such disruption or weakening of interactions could reverberate through the entire network and thereby trigger large irreversible changes. It is thus crucial to understand the dynamics of mutualistic networks and their responses to warming as well as the potential for the emergence of critical transitions that could impact loss of biodiversity.

The mutually beneficial interactions between plants and pollinators have been an important facet in the maintenance of global biodiversity. These interactions involve numerous species that co-evolve in response to various factors, including, but not limited to, changes in the environment, shifts in interaction dynamics, or both. Shifts in interaction strength could be due to differences in antagonistic interactions relative to mutualistic interactions (Andreazzi *et al*. 2020). Particularly, plant-pollinator interaction networks are increasingly endangered by both local or global extinctions, primarily due to alterations in the external environment caused by human activity (Biesmeijer *et al*. 2006, Memmott *et al*. 2007). For instance, due to anthropogenic climate change, there could be a disruption of seasonal timing of flower production or pollinator activity leading to reduced interactions between plants and animals (Harrison 2000, Wall *et al*. 2003, Memmott *et al*. 2007). Consequently, the rising temperatures caused by anthropogenic activities have the potential to impact the phenology of both plants and pollinators. In response to temperature increases, plants have been observed to flower earlier, directly indicating their vulnerability to temperature changes (Miller-Rushing *et al*. 2008, Miller-Rushing & Primack 2008, Menzel *et al*. 2006, Hegland *et al*. 2009). More often, insect-pollinated plants are particularly susceptible to climate warming since they depend on pollinators for their emergence, and the developmental life cycles of most pollinators are directly influenced by temperature. One of the major pollinators, Apis mellifera L, showed a relationship between spring temperatures and its first emergence (Gordo & Sanz 2005). Given that the phenology of both plants and pollinators relies on temperature, any alterations or increases in temperature will inevitably impact plant-pollinator interactions and potentially impact the stability and resilience of these communities.

Complex ecological systems, including but not limited to plant-pollinator systems, exhibit alternative stable states (Lever *et al*. 2014, Baruah 2022, Rohr *et al*. 2014). Such systems can suddenly transition to a new alternative undesirable state as environmental conditions deteriorate. Mutualistic plant-pollinator networks do exhibit such sudden transitions which are interdependent on the architecture of species interactions. For instance, mutualistic network architecture defined by nestedness or connectance has significant impacts not only on the maintenance of biodiversity (Bastolla *et al*. 2009) in such networks but also on abrupt collapses and timing of such network collapses Baruah (2022), Lever *et al*. (2014). Given that such networks exhibit alternative stable states and therefore could suddenly tip over from a desirable functional state to an undesirable state due to small changes in environmental conditions, it is thus crucial to understand how increases in temperature could potentially cascade through such networks and impact stability biodiversity, and whether such networks could potentially adapt to such changes.

Recent studies have reiterated on understanding the maintenance and resilience of biodiversity from an eco-evolutionary perspective (Dakos *et al*. 2019, Matthews *et al*. 2011, Norberg *et al*. 2012, Åkesson *et al*. 2021, Lasky 2019). In this context, trait variation plays an important role and acts as an engine in influencing biodiversity patterns as well as on the resilience of ecological communities (Barabas & D’Andrea 2016). Trait variation could both negatively or positively impact species diversity depending on whether the interaction among species is positive or negative (Baruah 2022, Baruah *et al*. 2022a). In addition, the degree of trait variation either facilitates the onset of a tipping point or delays it (Dakos *et al*. 2019, Chaparro Pedraza *et al*. 2021). Such trait variation is widespread among species embedded in such a complex web of interaction networks. For instance, in the context of mutualistic plant-pollinator interactions, such intraspecific variation could arise when individuals differ in their phenology, i.e., in their emergence or onset of flowering for plants that depend concurrently on temperature (Hegland *et al*. 2009). Such variation in phenological responses would therefore impact species interactions as increases in temperatures due to anthropogenic changes occur. Thus far, eco-evolutionary changes in phenological traits of plants and pollinators in response to warming need to be understood in order to forecast future unwanted changes to mutualistic communities.

In this study, we aimed to understand the eco-evolutionary response of mutualistic networks in response to warming using 101 empirical networks. We model a phenological trait of species that is involved in mutualistic interactions that is consequently also dependent on local environmental temperature. This phenological trait, the optimum temperature, also determines an individuals rate of growth. We show that eco-evolutionary feedbacks as temperature rises determine the resilience of mutualistic plant-pollinator networks. Variation in trait and evolutionary dynamics of the phenological trait in response to warming significantly increase the resilience of mutualistic networks and delay the occurence of tipping points. In addition, presence of eco-evolutionary feedbacks further decreases the occurrence of hysteresis in these networks. Our results point towards the fact that the architecture of such networks is a evolutionary spandrel that concurrently interact with eco-evolutionary dynamics to determine resilience in response to a rise in temperature.

## 2 Methods: modelling framework

We model a phenological trait *z* as a primary quantitative optimum trait that determines the intrinsic growth rates of species as well as trait-dependent interactions. This quantitative trait is normally distributed with mean *µ_i_* and variance of 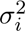. The per-capita growth rate of a phenotype *z* due to temperature-dependent growth, inter- and intraspecific competition, and mutualistic plant-pollinator interactions can be written as (shown here for the pollinator) (Baruah 2022, Valdovinos 2019):

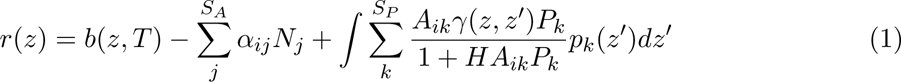

where *b*(*z, T*) is the temperature-dependent rate of growth, or tolerance to local environmental temperature T, *N_j_* is the pollinator density, *α_ij_* is the matrix of inter- and intra-specific competition, *A_ik_* is the element of the adjacency matrix denoting either 0 or 1, where 0 indicates no interaction and 1 indicates that species *i* interacts with species *j*, *H* is the handling time and *p_k_*(*z^′^*) is the distribution of trait *z^′^* which we consider to be normally distributed. *γ*(*z, z^′^*) is the trait-based interaction kernel of plants and pollinators which is also dependent on their optimum traits *z* and *z^′^*. If two individuals belonging to either a plant species or an animal species have similar optimum temperature traits, this would indicate that their chances of interaction are high given by a Gaussian interaction kernel as

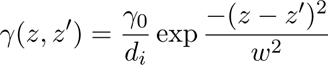

*γ*_0_ is the average interaction strength when a pollinator individual with optimum *z* interacts with a plant with optimum trait *z^′^*; *d_i_* is the degree of connectance of species *i*. In mutualistic interaction networks, generalist species generally benefit from a disproportionate number of interactions. To account for such asymmetries in mutualistic interactions, we used a trade-off that takes into account the number of interactions and average mutualistic strength given by 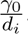. This formulation indicates that the strength of mutualistic interaction falls off as the degree of species increases (Dakos & Bascompte 2014, Lever *et al*. 2014, Baruah 2022) ensuring that generalist does not enjoy disproportionate densities. Empirically in nature, this value could be between 0 and 1 (Simmons *et al*. 2020). Temperature-dependent growth or the tolerance curve follows from a modified version of a previous study of Amarasekare & Johnson (2017) using a Gaussian form as,

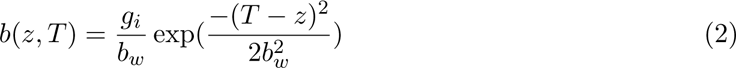

Here, *T* is the local temperature, *b_w_* and *g_i_* are the parameters that modulate the shape of the temperature tolerance curve. The denominator of this function, i.e. *b_w_*, modulates the width of the curve, and the ratio 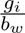 controls the peak of the temperature tolerance curve (Amarasekare & Johnson 2017, Åkesson *et al*. 2021). With this the population dynamics of species *i* and evolutionary trait dynamics of the mean trait *u_i_* can be written as:

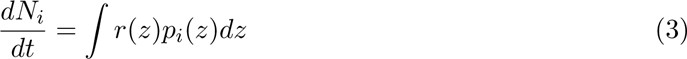

and,

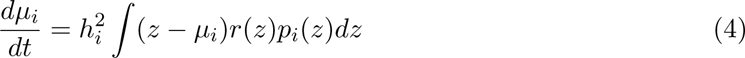

Equations 3 and 4 describe the ecological and evolutionary dynamics of mutualistic plant-pollinator networks in response to selection pressures arising either due to changes in local temperature *T* or indirectly due to shifts in interaction strengths. Note that the index *i* denotes a pollinator species *i*, and similar equations follow for plant species too (see supplementary 1). 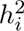 is the broad-sense heritability of the mean trait *µ_i_*.

### 2.1 Evaluating the temperature at which network collapses, and chances of abrupt collapses

To model the eco-evolutionary dynamics of plant-pollinator networks, we extracted 86 plant-pollinator networks from www.web-of-life.es database. In our modeling framework, we made the assumption that competition is generally weaker compared to average mutualistic strength, *γ*_0_. *γ*_0_ was fixed at 1.5 (see details in table 1). Additionally, intraspecific competition was set at a fixed value of 1, which was assumed to be much stronger than interspecific competition (Barabás *et al*. 2017, Baruah 2022). Interspecific competition was randomly selected from a uniform distribution ranging from *U* [0.0001, 0.001].

**Table 1:** Parameters and their values with the description used in the study. *U* [*a, b*] is the uniform distribution between *a* and *b*.

| Symbol | Description |
| --- | --- |
| $\sigma_i$ | Trait standard deviation of species $i$; sampled from either $U[0.002, 0.009]$ (low), or $U[0.02, 0.09]$ (high). |
| $h_i^2$ | Heritability of species $i$'s trait; either 0 or 0.4. |
| $g_i$ | modulates the peak of temperature-dependent growth rate which was fixed at 1 for all species. |
| $b_w$ | modulates the trade-off between temperature-dependent growth rate width, and peak, which was fixed at 2 for all species. |
| $\alpha_{ij}$ | pairwise inter- and intraspecific competition. Intraspecific competition $\alpha_{ii}$ was fixed at 1. Interspecific competition $\alpha_{ij}$ was randomly sampled from $U[0.001, 0.005]$. |
| $d_i$ | degree of species $i$. |
| $\gamma_0$ | average mutualistic interaction strength which was fixed at 1.5. |
| $T$ | Average local environmental temperature. The values of local average temperature varied for each simulation from $10^\circ C$ to $42^\circ C$. |
| $\mu_i$ | Mean phenological trait values of species. Their initial values are sampled from $U[10, 32]$. |
| $N_i N_p$ | Population density of animal and plant species. Their initial value was 1 for all species. |

Our next step was to assess at which local temperature *T* mutualistic plant-pollinator networks collapsed for different levels of trait variation (high versus low) in the mean optimum trait. Additionally, we examined the effect of evolutionary dynamics, i.e. heritability *h*^2^ either was 0.4 or 0, indicating that the mean phenological trait variation was either heritable and thus evolvable, or not heritable and therefore not evolvable. To evaluate the temperature at which the studied plant-pollinator networks collapsed, the local temperature was gradually increased from 10*^◦^C* to as high as 42*^◦^C* in steps of 0.5*^◦^C*, and equilibrium eco-evolutionary dynamics of 86 networks were assessed. The initial starting density of each species in each mutualistic network was fixed at 1 and the mean optimum trait *µ_i_* for each species in a mutualistic network was sampled from a uniform distribution of *U* [10, 32] (see table 1 for parameter values and description). For high variation in mean trait *µ_i_* we sampled trait variance from a uniform distribution of *U* [0.02, 0.09] and for low trait variation we sampled from 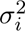 from *U* [0.002, 0.009]. For each different treatment of trait variation, the heritability of the mean trait and local environment temperature *T*, we simulated eco-evolutionary dynamics (equation 3, 4) for a duration of 10^4^ time points. We tracked species mean trait values and species densities over time and calculated network richness, and biomass of networks over time. We classified a species in a network to be extinct if its density fell below 0.01.

Next, for each species in the network, we evaluated whether the temperature at collapse and occurrence of abrupt collapses depended on network architecture, heritability of the phenological trait (*h*^2^ = 0*, h*^2^ = 0.4), and trait variation. We classified a network to be collapsed if the total equilibrium density (sum of the densities of all species in a network at the final time point of *t* = 10^4^) of the network fell below 0.1 for any local temperature T.

Furthermore, we defined whether a network displayed abrupt collapses as follows: for each local temperature T, we also computed the equilibrium network richness, which is the richness of the plant-pollinator community at the final time point of 10^4^. If the difference in equilibrium network richness at the temperature at the point of collapse and the equilibrium network richness at the preceding temperatures exceeded 10% of the total network size, we classified the network as exhibiting abrupt collapses. Conversely, if the difference in equilibrium network richness between the temperature at collapse and the penultimate temperatures was less than 10% of the network size, we categorized the network as not exhibiting abrupt collapses. In this way, we evaluated whether trait variation, evolutionary dynamics and network architecture interacted to impact the occurrence of abrupt collapses (Baruah 2022).

### 2.2 Stability at equilibrium of mutualistic networks

At the end of our simulations for each temperature, for each level of trait variation and heritability, we measured the (ecological) stability of each plant-pollinator network by first calculating the community matrix (Jacobian of the ecological part of the dynamics, evaluated at quasi-equilibrium; Supplement2) and obtaining its dominant eigenvalue. The dominant eigenvalue denotes the rate at which a given perturbation if applied to an ecological system at equilibrium will either contract or expand. If the dominant eigenvalue is negative, the perturbation will decay at the rate given by the absolute real value of the dominant eigenvalue and thus will indicate that the community is stable.

The Jacobian of the ecological part of the system at equilibrium is given by (see detail derivation in the supplementary 2):

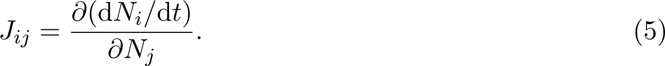

### 2.3 Capacity of networks to adapt: network trait-lag

Additionally, we estimated the capacity of each mutualistic network to adapt to changes in local temperature by calculating community trait lag as the lag between the mean community trait and local environment at the end of each simulation. This we did for each combination of the two levels of heritability, the two levels of trait variation and environment temperature. (as changes in local temperature at the end of each simulation for two levels of heritability, two levels of trait variation.) We thus define the mean trait lag of each mutualistic network as:

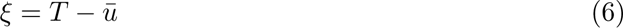

where *T* is the local temperature, and 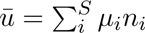, where *n_i_* is the relative species density at the end of each simulation run. In addition, we also quantified trait matching (see supplementary 1 for details) in networks as a measure of trait overlap between species in a network.

## 3 Results

We modelled the eco-evolutionary dynamics of 86 plant-pollinator networks in response to rise in temperature 1). Network size in the 86 plant-pollinator networks ranged from as low as 16 to as high as 104. Network connectance ranged from a low of 0.066 to a high of 0.42, and nestedness (NODF) ranged from a low of 0.111 to a high of 0.85.

### 3.1 impact of trait variation and heritability on temperature at collapse

When the heritability of the phenological trait was zero, which meant the absence of evolutionary dynamics, mutualistic networks collapsed at a much lower temperature (mean of 29^0^*C*, s.e = 0.70) than when the heritability of the phenological trait was high (mean of 33^0^*C*, s.e = 0.26). In the absence of evolutionary dynamics i.e., *h*^2^ = 0, whether species exhibited high or low trait variation did not play a role with respect to the temperature at which networks collapsed i.e., in the case of 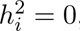, networks on average collapsed at similar temperatures for both high or low trait variation (mean of 29.2^0^*C*, s.e = 0.70 for low trait variation and mean of 28.5^0^*C*, s.e. = 0.85 for high trait variation. However, in the presence of evolutionary dynamics, i.e. *h*^2^ = 0.4, whether species exhibited high or low trait variation played a role concerning the temperature at which the networks collapsed. On average, networks with high trait variation collapsed at around 34.1^0^*C*, two degrees higher than when networks had low trait variation which was around 32.3^0^*C*.

### 3.2 Impact of network architecture on the temperature at collapse and occurrence of abrupt collapses

Our results indicate that network architecture interacts with evolutionary dynamics to impact the temperature at which networks collapse. Larger networks collapsed at higher temperatures than smaller networks (Fig 3A) for both when species had high or low trait variation. This was more prominent when heritability was 0.4 and when species exhibited high trait variation (Fig. 3A). The relationship between temperature at collapse of plant-pollinator networks and network connectance was overall negative. On average networks with higher connectance collapsed at lower temperatures than networks with low connectance. The slope of this relationship was not different between low or high trait variation when heritability was zero, or when heritability was positive. The presence of evolutionary dynamics delayed the temperature 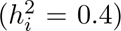 at which networks collapsed more so in the case of high trait variation and when networks had low connectance than when networks at high connectance (Fig. 3B). With nestedness (NODF), we see a similar relationship as we observe with network connectance (Fig. S2).

In addition, we analyzed how abrupt collapses were related to network architecture such as the size of the network, connectance, and nestedness (Fig. S2). Chances of abrupt collapses decreased as network size increased irrespective of whether species had high or low trait variation or whether heritability was zero 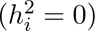 or positive 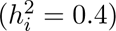 (Fig 3C). There were little to no differences in terms of chances of abrupt collapse w.r.t different levels of trait variation or heritability. Furthermore, the chances of abrupt collapse increased as connectance or nestedness increased (Fig. S2). In the absence 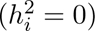 or presence of evolution 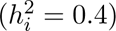, highly connected networks were more likely to exhibit abrupt collapses than lowly connected networks (Fig. 3D).

### 3.3 Local stability of networks: impact of warming, trait variation, heritability, and network architecture

Network architecture interacted with evolutionary dynamics to stabilize plant-pollinator networks. For instance, as network size increase, or nestedness (NODF) of networks increased, the dominant eigenvalue at equilibrium decreased more so when species had high trait variation and when evolutionary dynamics were at play (Fig. 4A). This was, however, not observed when heritability was off, or when species had low trait variation indicating that heritable variation impacted community stability, and made some networks more stable than others.

As warming increased, the dominant eigenvalue became more negative indicating that networks became more stable. The stability peaked at temperatures 17^0^*C* to around 28^0^*C* (supplementary Fig. S3). However, as the temperature kept rising, the stability of networks at quasi-eco-evolutionary equilibrium decreased drastically. In addition, when evolutionary dynamics were at play i.e., heritability of the mean trait was positive, and with high trait variation, plant-pollinator networks were overall more stable than when species had low trait variation or when heritability was switched off (supplementary Fig. S3).

### 3.4 Capacity of networks to adapt to temperature change

Mean community trait lag that determines whether the mutualistic community was able to track the optimum local temperature and thus adapt was dependent on the presence of evolutionary dynamics as well as on levels of trait variation. Higher trait variation and higher heritability of mean phenological trait led to low trait lag as local environmental temperature increased (Fig 4B). However, when heritability was zero, the trait lag substantially increased at lower local environmental temperatures than when heritability was high for high or low trait variation.

There was a shift in the trait lag at high temperatures for both levels of heritability as well as trait variation which marked the onset of community collapses. Network trait lag still kept increasing at very high temperatures as species in these networks were unable to evolve to track the changes in the temperatures as species densities remained near zero or very low (Fig. 4B).

## 4 Discussion

Changing environmental conditions such as warming pose a threat to the maintenance of biodiversity (Memmott *et al*. 2007, Ibáñez *et al*. 2010, Hegland *et al*. 2009, Kratina *et al*. 2012, Menzel *et al*. 2006). Species interactions, survival, and movement ecology could potentially be impacted by rising temperatures. Such rise in temperature could also impact evolutionary processes including genetic adaptation to climate change (Amarasekare & Johnson 2017) or tracking the rise in temperature through phenology (Hegland *et al*. 2009, Biesmeijer *et al*. 2006, Menzel *et al*. 2006). However, despite recognizing the interplay between ecological and evolutionary processes in determining species’ survival and interactions under modified climate conditions, studies considering the combined effects of these processes are far and few (Åkesson *et al*. 2021, Sales *et al*. 2021, Faillace *et al*. 2021, Cotto *et al*. 2017), particularly in terms of resilience to the occurrence of tipping points. Without the interplay of eco-evolutionary processes, the structure of species interaction in mutualistic networks was suggested to impact the occurrence of tipping points in response to rising temperatures (Bhandary *et al*. 2023). However, such mutualistic interactions could potentially evolve in response to rising temperatures and thereby further impact the occurrence of network collapses.

We model the eco-evolutionary dynamics of mutualistic systems to rising environmental temperatures. Specifically, we model an optimum trait of species that determines the rate of intrinsic growth or tolerance to local temperature as well as its interaction with other species. For instance, empirical plant-pollinator studies have demonstrated that the emergence of insect pollinators and plant flowering is determined by local environmental variables like spring temperature (Hegland *et al*. 2009, Gordo & Sanz 2005). As such, any change in local temperature could potentially disrupt plant-pollinator interaction thereby impacting the resilience of plant-pollinator interactions. We model this optimum trait in our study, and mutualistic interaction between plants and pollinators is dependent on how close they are in terms of optimum trait value i.e., if plants and pollinators have a similar average optimum trait, successful mutualistic interaction will take place. This follows from the fact that plant flowering temperature and insect emergence temperature should match for mutualistic benefits to occur (Hegland *et al*. 2009). Thus, the rate of growth of species was not only dependent on local temperature but also on whether their optimum trait matched with other species. This optimum trait of species thus evolved in response to selection pressures that arose from species interactions as well as due to a rise in temperature.

We found that at equilibrium, mutualistic network biomass reached higher values when species exhibited higher trait variation in contrast to when species exhibited lower trait variation (Fig.1-2A). When species had high trait variation, mean trait matching was much higher among plants and pollinators (Fig. S1). As a result, on average, strength of mutualistic interaction increased. Consequently, fitness benefits in terms of increases in the rate of growth were higher leading to higher density when species had higher trait variation, in comparison to when species had low trait variation (Fig. 1, Fig. 2). In addition, in the absence of evolution, that is, when heritable variation was zero 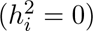, networks had lower richness and lower biomass in contrast to when the trait was heritable. Naturally, due to heritable trait variation, species were not only able to adapt to the local environmental temperature but also had higher trait matching in general (Fig. S1). This resulted, on average, higher richness and higher biomass particularly in the case when species had high trait variation. At low levels of heritable trait variation i.e., in the case when 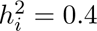 and *σ_i_ ɛ U* [0.002, 0.009], evolutionary dynamics were hampered. Faster evolutionary dynamics as well as higher trait matching of plant and pollinator optimum traits would occur only when there was a sufficient amount of variation in the mean trait value(Yoshida *et al*. 2003). Consequently, even when heritability was high, low trait variation would still lead to lower adaptation not only to local environmental temperature but would also impact the average mutualistic strength of interaction as there would be lower trait overlap, in comparison to when species exhibited high trait variation (Fig. S1). Thus, trait variation as well as heritability of the optimum trait would significantly influence whether networks would be able to have a higher functionality in the face of climate change.

**Figure 1:**
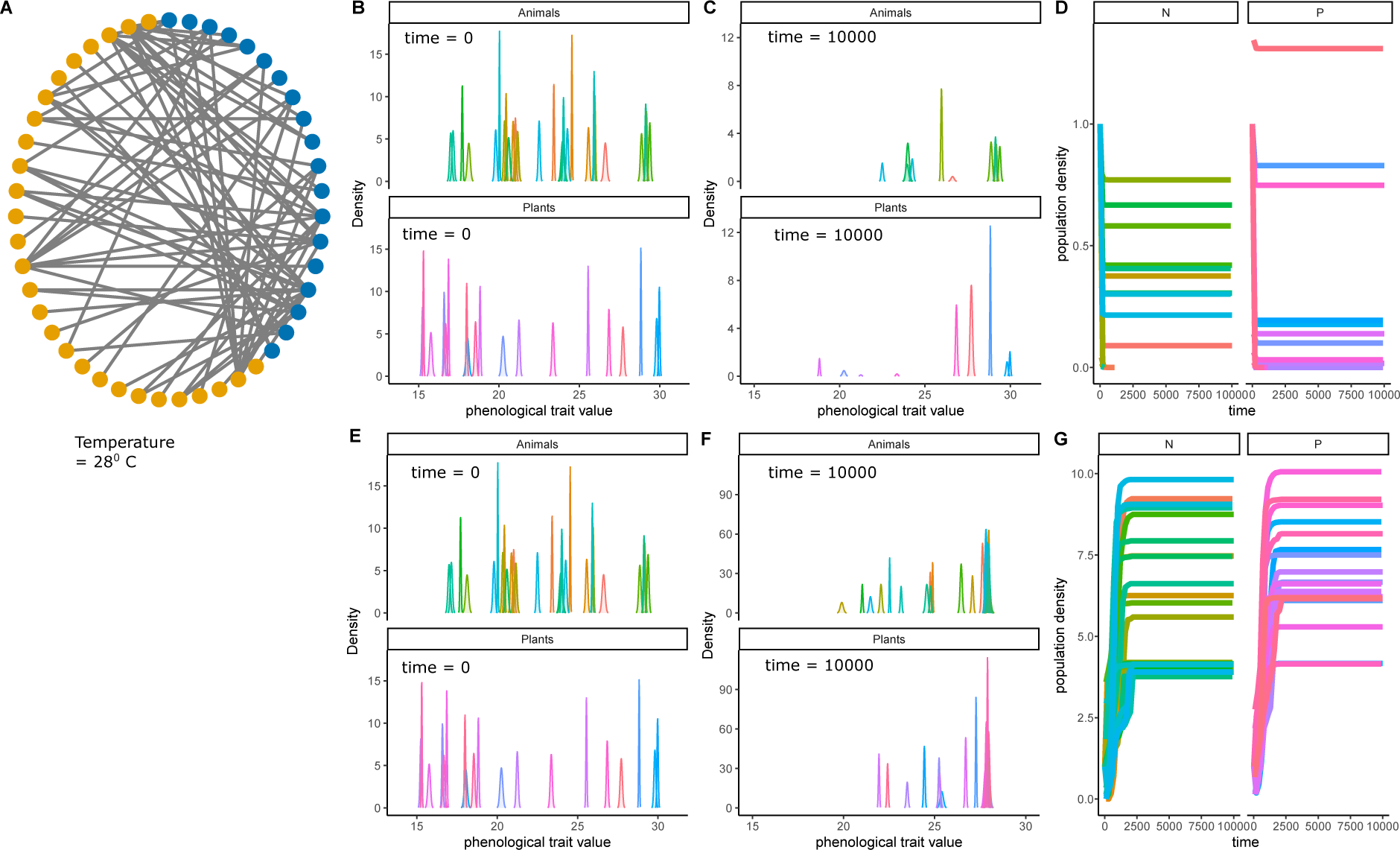
The effect of warming on equilibrium eco-evolutionary dynamics of plant-pollinator networks. A) An example forty-seven species plant-pollinator network. (B) The initial density of the optimal phenological traits of plants and animals, for a local environmental temperature of 28*^◦^C*. (C) At *t* = 1*e*^4^, the optimum phenological trait density of the surviving species clustering around 28*^◦^C* which was the local environmental temperature. (D) Population density of plants and animals over time. (B-D) In these three plots, heritability was zero, i.e. the trait was not heritable, 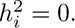. This means evolutionary dynamics were not at play. (E-F) Phenological trait density at time point *t* = 1 and at time point *t* = 1*e*^4^. Note that trait density at evolutionary equilibrium was much higher than in (C). Also note that in E-F, heritability was high, i.e. 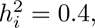, for all species, and thus evolutionary dynamics of the optimal phenological trait were in play. (G) Population density of plants and animals over time. In all the simulations, initial trait variance was randomly sampled from *U* [0.02, 0.09].

**Figure 2:**
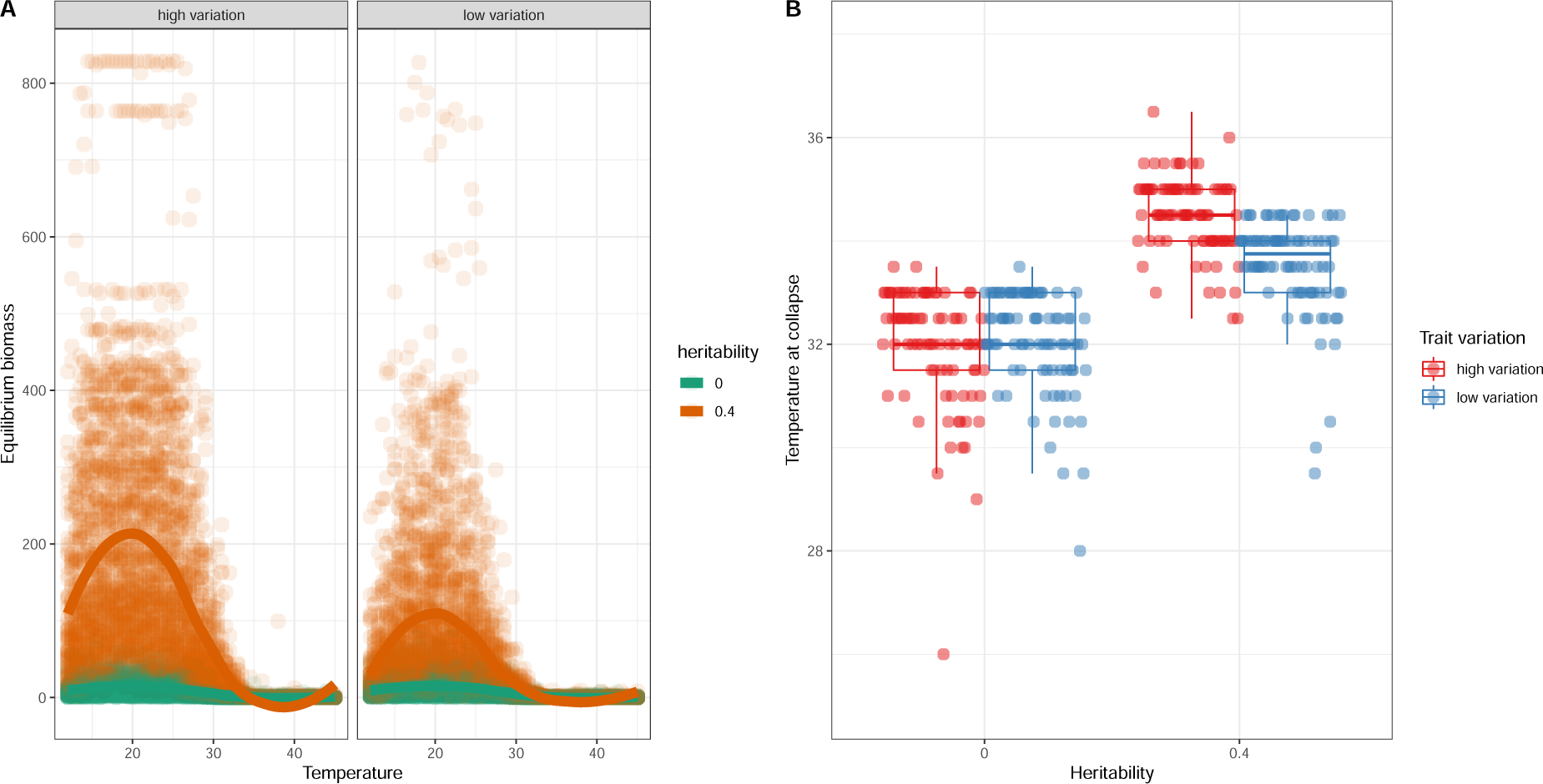
Temperature at collapse for 86 mutualistic networks for two levels of trait variation and two levels of phenological trait heritability. A) Total network biomass of mutualistic networks as environmental temperature increased for two levels of trait variation (high and low) and two levels of heritability (0 and 0.4). Each circle is the equilibrium network biomass at the final time point *t* = 10^4^ for a particular temperature. B) Boxplots of temperature at collapse for all 86 networks analyzed for two levels of trait variation (high or low) and two levels of heritability (0 and 0.4).

The resilience of mutualistic networks has been subject to much scrutiny (Bastolla *et al*. 2009, Lever *et al*. 2020, 2014, Dakos & Bascompte 2014, Jiang *et al*. 2018, Bascompte & Stouffer 2009, Baruah 2022, Arroyo-Correa *et al*. 2023, Baruah *et al*. 2022b). Mutualistic networks generally exhibit alternative stable states and show sudden collapses as the strength of mutualistic interaction decreases (Baruah 2022, Lever *et al*. 2014, Jiang *et al*. 2018) or when the loss of species occurs (Fortuna & Bascompte 2006). Previously, when such mutualistic plant-pollinator networks exhibit some amount of trait variation, abrupt collapses occur more often than not. Although abrupt collapses are more often observed when networks have a higher amount of trait variation, the threshold at which such networks usually collapsed did not differ much when compared between high or low levels of species trait variation (Baruah 2022). In our study, species growth rates and the evolution of the mean optimum trait were impacted through two avenues. One was the selection pressure to adapt to the local environmental temperature, and the other was the selection pressure to match the partner’s phenological optimum temperature for successful mutualistic positive interaction. As such, when the local environmental temperature was within the optimum phenological temperature range of species being sampled, network richness and biomass were high, especially when species had high heritable trait variation. As the local environmental temperature was outside the range of the optimum traits of species i.e., outside 30*^◦^C*, networks collapsed. However, the temperature at which networks collapsed significantly differed when species had high heritable trait variation or not. Particularly, the heritability of the mean trait and the presence of high trait variation in species’ mean optimum trait delayed network collapses than when species had low trait variation or the optimum phenological trait was not heritable. Our results suggested that evolutionary dynamics and the presence of trait variation could delay the collapse of networks (Dakos *et al*. 2019). In the previous study of Baruah (2022), trait variation didn’t delay the occurrence of collapses in mutualistic networks. The difference is that in our study local environmental temperature not only impacted the mutualistic strength of interaction among species but also impacted the intrinsic rate of tolerance to the temperature. As a result, species in our study could evolve to have higher tolerance as local environmental temperature increased. In contrast, in Baruah (2022), the parameter that caused the networks to collapse only impacted the mutualistic strength of interaction and not the intrinsic rate of growth.

Nestedness is a property very commonly observed in mutualistic networks, where generalists interact with both specialists and generalists but specialist species interact only with generalists. Such a property enables a hierarchical structure of the community, with some species enjoying a disproportionate amount of mutualistic benefits than others. Such a network metric along with network connectance has been suggested to be important for stability as well as the resilience of networks to collapse (Bastolla *et al*. 2009, Lever *et al*. 2014, Baruah 2022). Previous studies have indicated that the architecture of mutualistic networks increases stability and resilience (Bastolla *et al*. 2009, Okuyama & Holland 2008, Thébault & Fontaine 2010, Baruah 2022). Our results further indicate that in the presence of evolution and high levels of trait variation, as nestedness increases or network size increases, local stability increases (Fig 4A, Fig. S6). With increases in nestedness, certain species i.e., the most generalist species are the ones that will have the most number of interactions. As a consequence, while assembling such a mutualistic network any specialist species entering such a community will enjoy the most mutualistic benefits and face less effective competition if it interacts with the generalists. This is due to the fact that generalist species have much higher abundances, which translates to higher mutualistic benefits for the specialist species. In such a way, a nested network architecture could promote higher biodiversity and stability (Bastolla *et al*. 2009). In addition, when such a mutualistic network exhibits high trait variation, specialist species abundance could further increase due to higher trait matching with generalist species (Baruah 2022). As a consequence, a nested network architecture not only would increase community biomass but also impact the stability of biodiversity. Our results corroborate this and indicate that network properties (nestedness, and network size) interacted with evolutionary dynamics to increase the local stability of mutualistic networks (Fig. 4A). Trait variation led to higher levels of trait-matching, and with higher levels of trait matching, species were able to maintain positive abundances even at high temperatures. As a result, local stability increased as evolutionary dynamics delayed network collapses.

In our study, network architecture interacted with evolutionary dynamics to mitigate the threshold temperature at which networks collapsed (Fig. 3A-B). The relationship between the size of the network and the temperature at which networks collapsed was positive. This meant that larger networks collapsed at higher temperatures in comparison to smaller networks (Fig. 3A). This result was modulated by trait variation and heritability, with higher heritable trait variation, there was delayed collapse of networks. So why did larger networks collapse at higher temperatures than smaller networks? This essentially was due to the simple fact that larger networks harbored more species than smaller networks, and as a result, maintained higher biomass than smaller networks even at very high temperatures (Fig. S5), and eventually delayed the collapse (Baruah 2023). As a consequence, species maintained higher density at higher local temperatures eventually delaying network collapse. On the other hand, since network connectance was negatively correlated with network size (Pearson correlation coefficient of −0.69) i.e., smaller networks show a higher connectance than larger networks, as a result, as network connectance increased, equilibrium biomass decreased (Fig. S5). Our results thus indicate that network architecture interacts with eco-evolutionary dynamics to delay the threshold temperature at which networks collapse.

**Figure 3:**
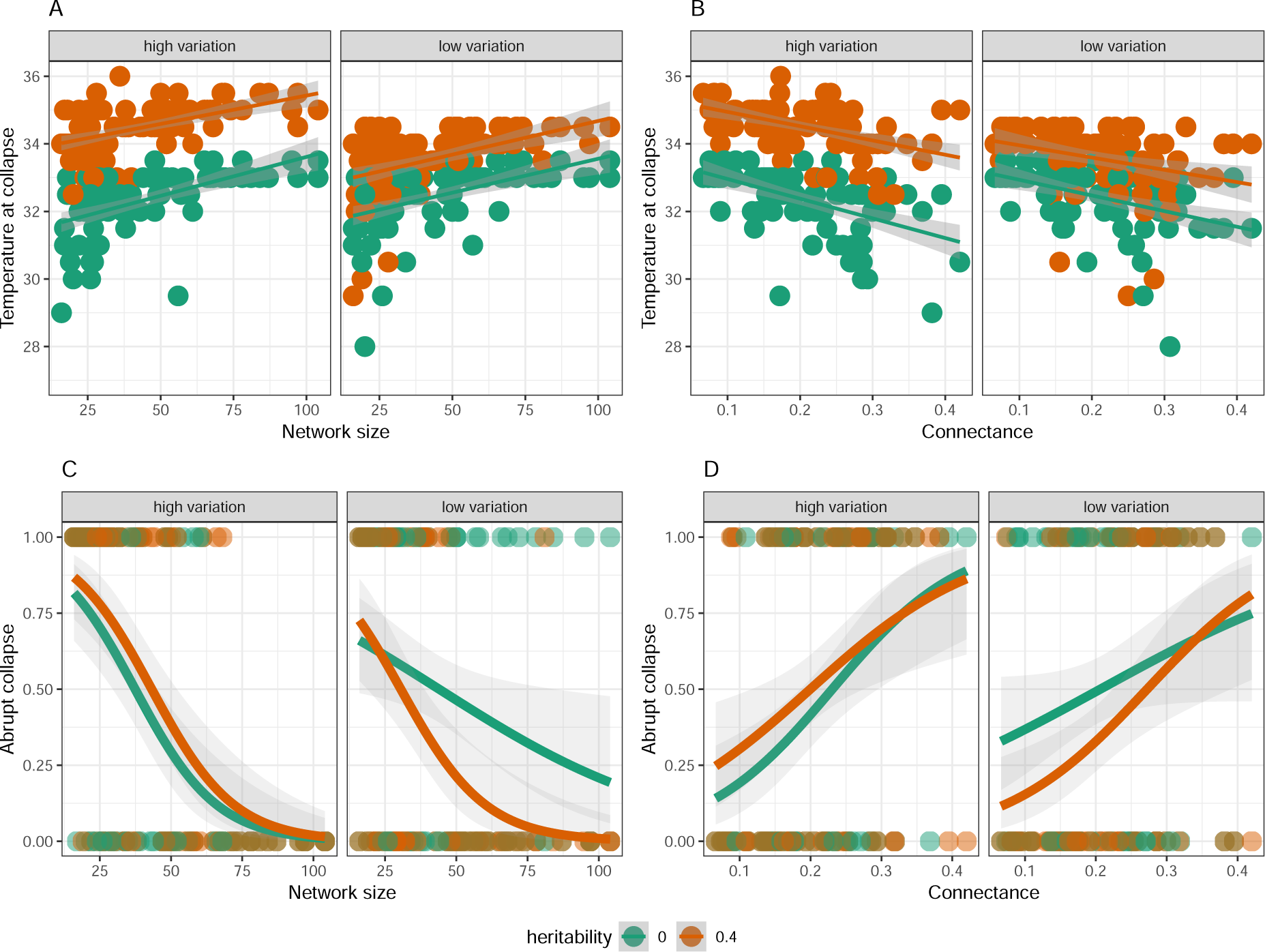
Temperature at collapse and chances for abrupt collapse in relation to structural properties of mutualistic networks. (A-B) The temperature at which networks collapsed was plotted in relation to network size (A) and connectance (B) for two levels of trait variation and two levels of heritability. (A) Smaller networks collapsed at lower temperatures in comparison to larger networks which was modulated by trait variation and heritability of the trait. This relationship was the opposite in terms of network connectance. In (A-B) lines represent linear regression model with standard error. (C-D) Chances of abrupt collapse for all mutualistic networks in relation to network size (C) and network connectance (D) for two levels of trait variation and two levels of trait heritability. Lines represent logistic regression with quasibinomial error distribution.

We demonstrate that in mutualistic communities, the lag in the average community trait’s response to local environmental temperature is contingent upon the degree of trait variation and whether the mean optimum trait is heritable (Fig. 4B). Greater levels of trait variation and heritability enable species to adapt to high temperatures. The synchronization of flowering or emergence between mutualistic partners is critical for the survival of both pollinators and plants. As the climate undergoes changes and environmental temperatures rise, alterations in the timing of flowering and emergence among pollinators, referred to as shifts in phenology, are inevitable, leading to widespread mismatches in their interactions (Hegland *et al*. 2009, Ibáñez *et al*. 2010, Memmott *et al*. 2007, Miller-Rushing & Primack 2008, Menz *et al*. 2011). These mismatches could result in reduced pollen deposition due to changes in visitation rates or insufficient pollination, potentially impacting population densities in plants (Miller-Rushing & Primack 2008). When heritable variation in the optimum trait was zero, mismatches occurred not only in terms of species interaction but as well as to local temperature. As a consequence, species could not adapt to either the rise in temperature or to species’ optimum traits for beneficial mutualistic interactions. Thus, community trait lag was considerably higher when variation in the mean optimum trait was low and heritability was zero. However, these mismatches could be reduced if both plants and pollinators had a sufficient amount of variation in their traits. In response to changes in the climate, fast evolutionary responses have been detected in flowering plants and insects (Davis et al 2005, Franks et al 2007). Indeed, a fast evolutionary response would depend on the amount of heritable variation in the optimum trait that we modeled. Indeed, a high amount of heritable variation in the mean optimum trait will provide a sustainable avenue for such species to adapt to increasing temperatures. However, if temperatures keep on rising, the tolerance to extreme temperatures itself needs to evolve for species to keep tracking changes in the temperature. In our model, since tolerance width is a fixed constant, consequently, at extreme temperatures, we do observe community trait lag to increase after local temperatures rose above 34*^◦^C*, the threshold at which community biomass of networks starts decreasing considerably.

**Figure 4:**
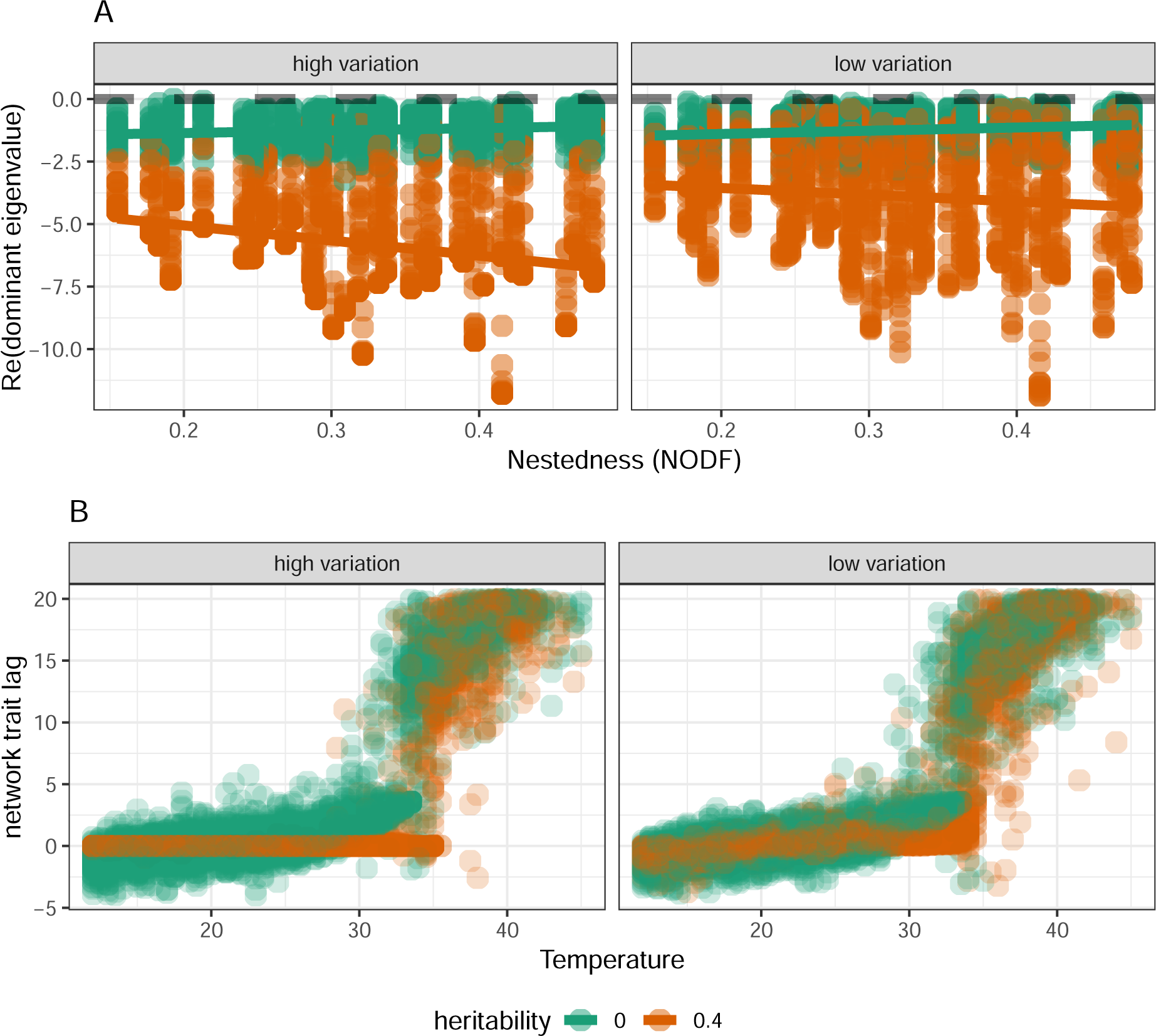
A) Local stability of 86 plant-pollinator networks in relation to network nestedness (NODF) and (B) ommunity trait lag of mutualistic networks in relation to increasing environmental temperature for two levels of trait variation and two levels of trait heritability. In (A) dominant eigenvalue was collated for local temperatures below 25^0^*C*. Lower values indicate more local stability of networks. (B) Each point is a trait lag estimated at equilibrium for a network. The dashed black horizontal line indicates when the mean community trait lag is zero. For high trait heritability 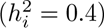 and high trait variation, trait lag was zero even at as high a temperature as 34*^◦^C*. Points above 35^0^*C* are the trait lag of the collapsed networks.

There are considerable variations in plant and pollinator life-history traits as well as thermal tolerance to warming. However, their responses to increasing temperatures are bound to be similar in plant-pollinator networks. As temperatures rise, mismatches in terms of phenological responses are bound to occur, and species might need to adapt their tolerance over time to maintain positive fitness. Our study points to the fact that indeed the ability to track local environmental temperatures considerably depends on the amount of heritable trait variation on the phenotypic optimum trait that responds to the rise in temperature. In addition, species in such networks can adapt beyond their optimum tolerance and delay the eventual community collapses dictated by a host of community properties such as community nestedness or size of the community. Our results indicate that the structural properties of a plant-pollinator network are intimately interlinked with how species can evolve and track changes in temperature to delay eventual network collapses. We argue that individual trait variation in association with the structural properties of a network plays a crucial role in mitigating changes in such networks in response to warming.

## Acknowledgments

GB would like to acknowledge DFG Walter Benjamin grant no BA 7974/1-1 for funding the research.

## Author contributions

GB conceptualised the study. TL did the simulations and analysed the results with GB. GB and TL wrote the manuscript.

