## appendix for "Mutualistic network architecture and eco-evolutionary feedbacks modulate the occurrence of transitions and stability in response to rising temperature"

### 1 General eco-evolutionary framework

We model the dynamics of pollinators N and plants P in an ecologically relevant quantitative trait $z$ that relates to its optimum temperature. Each individual belonging to a guild of pollinators or plants can be described with its trait $z$, but each of the species belonging to the guild of pollinators or plants is comprised of individuals with different trait values. Now the number of individuals with within species $i$ at time $t$ for pollinators will be $N_i(t)$ and for the plants it will be $P_i(t)$, and the distribution of their traits within each each species $i$ can be given by a function $p_i(z,t)$ and by definition this function satisfies

$$\int p_i(z,t)dz = 1$$

at every time $t$; the limits of integration encompass the whole trait axis, which for simplicity we take to go between minus and plus infinity. $N_i(t)p_i(z,t)dz$ is then the population density of species i's individuals with phenotype value between $z$ and $z+dz$ for the plants and for the pollinators, we can write $N_i(t)p_i(z',t)dz'$ which is the population density with phenotype value between $z'$ and $z'+dz$.

We work in the quantitative genetic limit, i.e., the optimum trait in question is determined by many independent loci (Bulmer 1980, Falconer 1981) and the trait distribution is normal and variance of the trait does not change in response to selection,

$$p_i(z,t) = \frac{1}{\sqrt{2\pi\sigma_i^2}} \exp \frac{-(z-u_i(t))^2}{2\sigma_i^2},$$

where $u_i(t)$ is the mean optimum temperature trait value for the species $i$ and $\sigma_i^2$ is the trait variance. In this scenario, only the mean of the trait responds to selection and the trait variance remains constant.

The governing dynamical equations of population dynamics can be written with Lotka-Volterra equations. The per-capita growth rate of pollinator, $r_N(z)$, and plants, $r_i^P$, can be written as:

$$r_N(z) = b(z,T) - \sum_j \alpha_{ij} N_j + \int \sum_k P_k \frac{\gamma(z,z')A_{ik}}{1+H\gamma(z,z')P_k} p_k(z',t)dz' \quad (1)$$

And,

$$r_P(z) = b(z,T) - \sum_j \alpha_{ij} P_j + \int \sum_k N_k \frac{\gamma(z,z')A_{ik}}{1+H\gamma(z,z')N_k} p_k(z',t)dz' \quad (2)$$

where $b(z,T)$ is the temperature-dependent trait-specific growth rate of phenotype $z$ independent of competition or mutualistic benefits for an animal and a plant (both have similar dependence). This means that an individual of a species belonging to a guild will be influenced by the local temperature $T$. H is the handling time. $A_ik$ is the adjacency matrix, and is 1 if species $i$ interacts with species $k$ and 0 if there is no interaction. $\alpha_{ij}$ is the pairwise competition term among species belonging to each own guild. We could however make this also trait-dependent by bringing a function that leads to trait-trait interaction. However, just for simplicity, we are going to assume that competition is not influenced by our trait of interest $z$; $\gamma(z,z')$ is the function that captures the mutualistic interactions among individuals. Here, $z$ is the optimum temperature of an individual of a species belonging to a guild say the pollinators, and $z'$ is the optimum temperature of an individual belonging to the plants. We can take this function to be a Gaussian:

$$\gamma(z,z') = \frac{\gamma_0}{d_i} \exp \frac{-(z-z')^2}{w^2}$$

where, $\gamma_0$ is the average strength of mutualistic interactions and $w$ is the width that controls how strongly two individuals interact. If the optimum temperature of the two individuals belonging to two different species namely plants and pollinators are similar, the stronger is the mutualistic benefit. Biologically this means that, if the optimum temperature for plants for flowering and the optimum temperature for insects to emerge match, then chances of a mutualistic interaction increase. For instance, one could imagine this specific mutualistic interaction occurring due to optimum temperature being similar for both individuals. Species phenology, i.e., flowering times of insect-pollinated plants are strongly dependent on temperature. Same goes for pollinator emergence which is also suggested to be correlated to monthly temperature. Thus, if optimum species phenotype i.e., plant and pollinator phenology matches then mutualistic interaction would be successful (Hegland et al 2009).

Equation 1 represents the per-capita growth rate of an individual with phenotype $z$ interacting facilitatively with another individual with phenotype $z'$ belonging to a species of another guild and $p_j(z',t)$ is the distribution of the trait $z'$. The integration goes over the entire trait space and is summed for all the species belonging to a guild. This formulation of the model is special in the sense that growth and mutualistic interactions only depend on the phenotype $z$ but not on species identity.

Now the population dynamics of species $i$ over all trait space $z$ can be written as (shown here for the animals $N$, same equation goes for the plants, $P$):

$$\frac{dN_i}{dt} = N_i(t) \int r_N(z) p_i(z,t) dz \tag{3}$$

Substituting equation 1 into 2 we get

Since the competition among species within a guild is independent of the phenotype we modeled and from $\int p_i(z,t)dz = 1$, we get,

$$\int \sum_j \alpha_{ij} N_j p_i(z,t) dz = \sum_j \alpha_{ij} N_j$$

$$\frac{dN_i}{dt} = N_i(t) \int \left( b(z,T) - \sum_j \alpha_{ij} N_j + \int \sum_k \frac{A_{ik}\gamma(z,z')P_k}{1+H\gamma(z,z')P_k} p_k(z',t)dz' \right) p_i(z,t)dz \tag{4}$$

$P_k$ is the density of plant species $k$. We can further solve equation 3 as:

$$\frac{dN_i}{dt} = N_i(t) \left( \bar{b_i} - \sum_j \alpha_{ij} N_j + \int \int \sum_k \frac{A_{ik}\gamma(z,z')P_k}{1+H\gamma(z,z')P_k} p_k(z',t) p_i(z,t) dz dz' \right) \tag{5}$$

where

$$\bar{b_i} = \int b(z,T) p_i(z,t) dz, \tag{6}$$

#### 1.1 Temperature, species interactions, and evolutionary trait dynamics

Following Amarasekare Johnson 2017, we model the temperature-dependence of $b_A(z,T)$ using the Gaussian form as (similar equation for the plants but shown here only for the animals):

$$b(z,T) = \frac{g_i}{b_w} \exp(-\frac{(T-z)^2}{2(b_w)^2}) - \kappa_i$$

where, $T$ is the current local temperature, $z$ is the phenotype of the individual that defines its optimum temperature, $\kappa_i$ is the extrinsic mortality; $a_w, b_w$ are positive constants that are used so that $b_A(z,T)$ mimic the growth rate as has been observed empirically in response to temperature. The width of this temperature tolerance curve is given by $2(b_w - a_w \mu_i)$ . This width parameter than causes a change in the temperature tolerance curve meaning that species with lower optimum mean phenotypic temperature trait, $\mu_i$ will have wider curves, whereas species with higher optimum trait, $\mu_i$ will have narrower curves. Integrating now equation 6 we get,

$$\bar{b_i} = \frac{g_i}{b_w}\exp(-\frac{(T-\mu_i)^2}{2b_w^2+2\sigma_i^2})\frac{b_w}{\sqrt{b_w^2+\sigma_i^2}} - \kappa_i$$

Evolutionary dynamics of the mean optimum temperature phenotype, or the mean phenological trait $\mu_i^A(t)$ of interest can then be written as and assuming in the quantitative genetic limit,

$$\frac{du_i}{dt} = h_i^2 \int (z-\mu_i) r(z) p_i(z,t) dz \tag{7}$$

where $h_i^2$ is the broad sense heritability, and $\sigma_i^2$ is the genetic variance which here is equal to phenotype variance.

we can further substitute from equation 4 into equation 5 as

$$\frac{du_i}{dt} = h_i^2 \int (z-\mu_i) b(z,T) p_i(z) dz - \int (z-\mu_i) \sum_j \alpha_{ij} N_j p_i(z) dz + \int\int \sum_k (z-\mu_i) \frac{\gamma(z,z')P_k}{1+H\gamma(z,z')P_k} p_k(z',t) dz' p_i(z) dz \tag{8}$$

here,

$$\int (z-\mu_i(t)) b(z,T) p_i(z) dz = \frac{g_i}{b_w}\exp(-\frac{(T-\mu_i)^2}{2b_w^2+2\sigma_i^2})\frac{\sigma_i^2 b_w (T-\mu_i)}{(b_w^2+\sigma_i^2)^{1.5}}$$

Similar, equation goes for the plant species except in the mutualistic interaction term where the species density would be replaced by $P$. Thus the two eco-evolutionary dynamical equations that would describe the dynamics of a mean quantitative trait $\mu_i(t)$ and dynamics of a species belonging to a guild, say the pollinators, are equation 5 and equation 8.

### 2 Jacobian matrix at equilibrium and stability of plant-pollinator networks:

The Jacobian of the ecological part of the system is given by

$$J_{ij} = \frac{\partial(\mathrm{d}N_i/\mathrm{dt})}{\partial N_j}. \tag{9}$$

We can write the ecological dynamics from either Eq. 5 as

$$\frac{dN_i}{dt} = N_i(t)\left(\bar{b_i} - \sum_j \alpha_{ij} N_j + \int\int \sum_k \frac{A_{ik}\gamma(z,z')P_k}{1+H\gamma(z,z')P_k} p_k(z',t) p_i(z,t) dz dz'\right) \tag{10}$$

We assume that the mutualistic system described above has a feasible equilibrium where the density of all species are positive, and therefore the equilibrium condition can be written as:

$$\bar{b_i} - \sum_j \alpha_{ij} N_j + \int\int \sum_k \frac{A_{ik}\gamma(z,z')P_k}{1+H\gamma(z,z')P_k} p_k(z',t) p_i(z,t) dz dz' = 0, \tag{11}$$

where $N_j$, $P_k$ are densities of animal and plants at equilibrium. The local stability at such an equilibrium is described according to whether the system returns to its equilibrium state following an infinitesimally small perturbation, and is thereby determined by the eigenvalues of the Jacobian matrix at equilibrium $J$. A sufficient condition for an equilibrium to be locally stable is if all the eigenvalues of the J has a negative real part. The real part of the dominant eigenvalue of the Jacobian determines the rate at which the perturbation expands or contracts. In order words, the real part of the dominant eigenvalue also represented as $Re(\lambda_1)$, determines the rate at which the mutualistic could recover. The more negative the value is the more stable the system is to a perturbation at equilibrium.

The diagonal elements of the Jacobian matrix at equilibrium can be calculated as:

$$\frac{\partial(\mathrm{d}N_i/\mathrm{dt})}{\partial N_i} = -\alpha_{ii} N_i \tag{12}$$

or,

$$\frac{\partial(\mathrm{d}P_i/\mathrm{dt})}{\partial P_i} = -\alpha_{ii} P_i \tag{13}$$

The off-diagonal elements of the Jacobian matrix can be calculated based on whether they represent competitive interactions or mutualistic interactions. If they represent competitive interactions they can be calculated as:

$$\frac{\partial(\mathrm{d}N_i/\mathrm{d}t)}{\partial N_j} = -\alpha_{ij} N_i \tag{14}$$

or,

$$\frac{\partial(\mathrm{d}P_i/\mathrm{d}t)}{\partial P_j} = -\alpha_{ij} P_i \tag{15}$$

And if they represent mutualistic interactions, it can be calculated as:

$$\begin{aligned}\frac{\partial(\mathrm{d}N_i/\mathrm{d}t)}{\partial P_k} &= \frac{\partial}{\partial P_k}(\int\int \sum_k \frac{A_{ik}\gamma(z,z')P_k}{1+H\gamma(z,z')P_k} p_k(z',t)p_i(z,t)dzdz') \\ &= (\int\int \frac{A_{ik}\gamma(z,z')N_i}{(1+H\sum_k \gamma(z,z')P_k)^2} p_k(z',t)p_i(z,t)dzdz')\end{aligned} \tag{16}$$

Or,

$$\begin{aligned}\frac{\partial(\mathrm{d}P_i/\mathrm{d}t)}{\partial N_k} &= \frac{\partial}{\partial N_k}(\int\int \sum_k \frac{A_{ik}\gamma(z,z')N_k}{1+H\gamma(z,z')N_k} p_k(z',t)p_i(z,t)dzdz') \\ &= (\int\int \frac{A_{ik}\gamma(z,z')P_i}{(1+H\sum_k \gamma(z,z')N_k)^2} p_k(z',t)p_i(z,t)dzdz')\end{aligned} \tag{17}$$

Or,

Thus, the Jacobian matrix at equilibrium can be written as,

$$J = J_1 + J_2$$

where,

$$J_1 = \begin{pmatrix} J_{cA} & 0 \\ 0 & J_{cP} \end{pmatrix},$$

where, $J_{cA}$ and $J_{cP}$ are the matrices that contains the elements competition from equation 12, 13, 14 and 15. And finally,

$$J_2 = \begin{pmatrix} 0 & J_{mA} \\ J_{mP} & 0 \end{pmatrix},$$

where, $J_{mA}$ and $J_{mP}$ are the matrices that contains the elements competition from equation **??**, 16, 17.

Thus the real part of the dominant eigenvalue can be calculated from the the Jacobian matrix at equilibrium , i.e., from J.

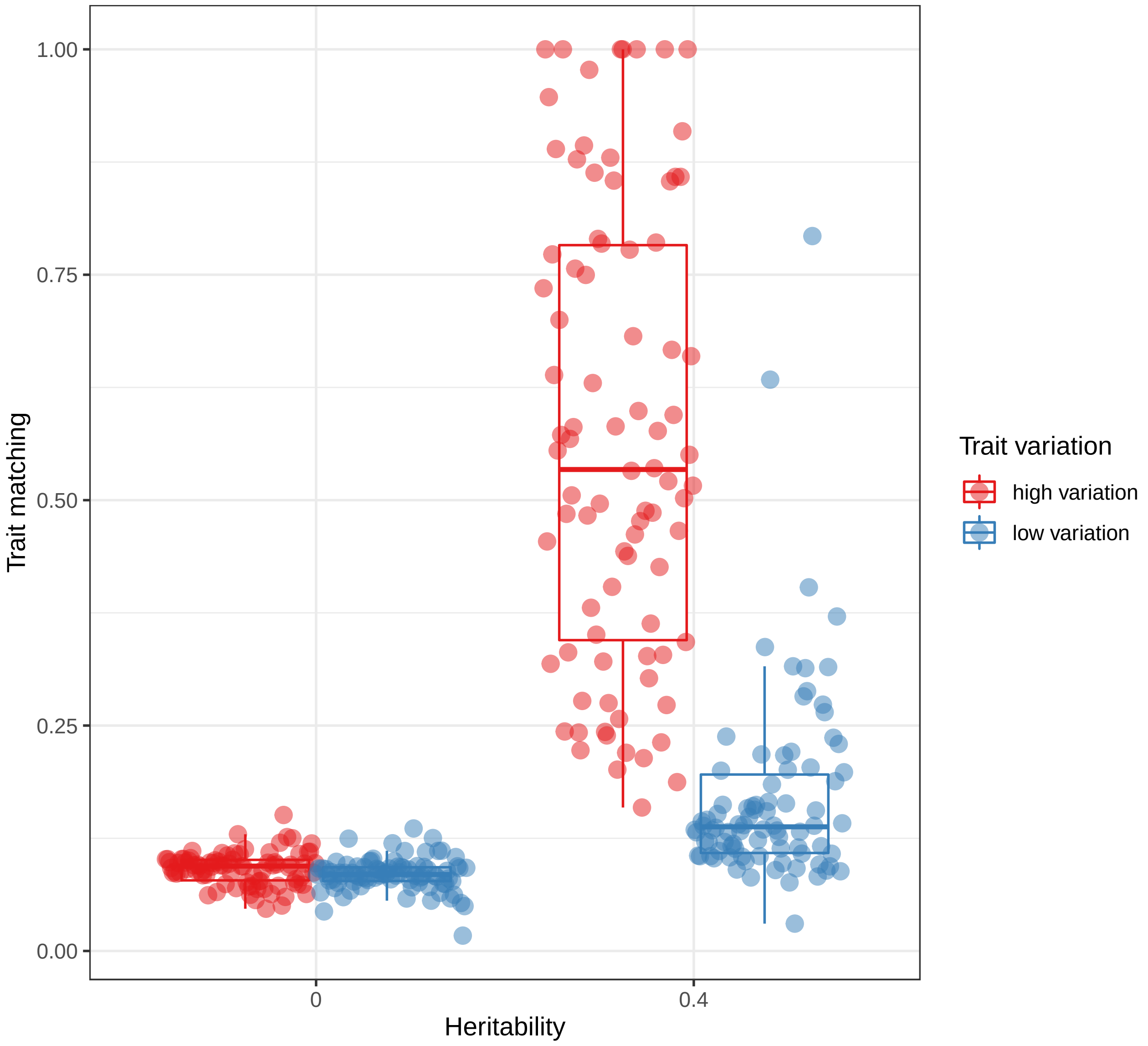


Figure S1: Mean network trait matching for 89 plant-pollinator networks for two levels of heritability and two levels of trait variation. Here trait matching between two species in a network was calculated as a metric for trait-overlap as: $\frac{\exp(-(\mu_i-\mu_j)^2)}{\sigma_i^2+\sigma_j^2}$. Then mean network trait-matching was calculated as mean over all species in a network. Here mean trait-matching can go from 0 to 1, where 1 is the max overlap between plant and pollinator mean traits. In the presence of evolution, trait matching was highest when species had high trait variation and lowest when evolution was absent.

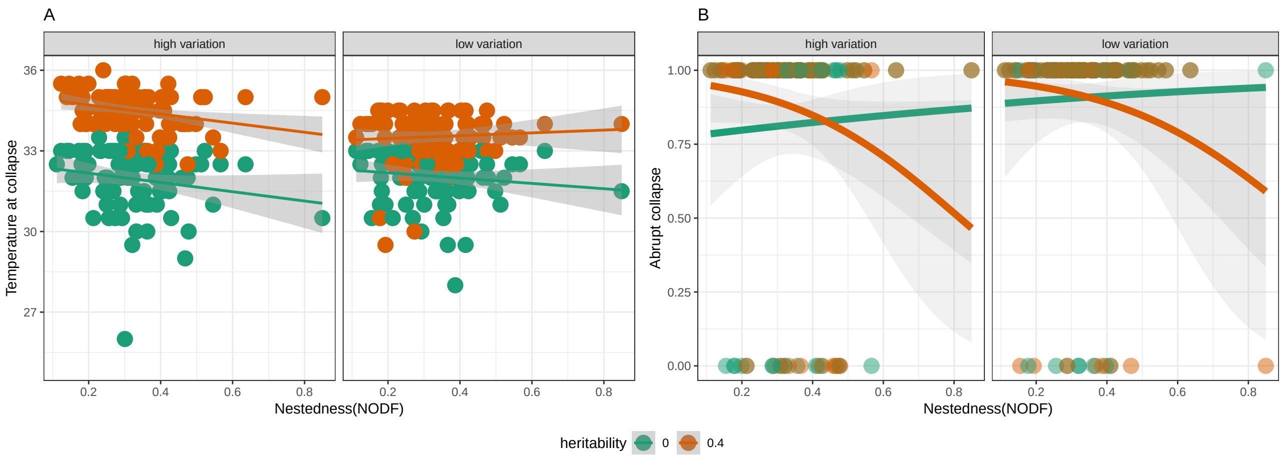


Figure S2: Temperature at collapse and chances for abrupt collapse in relation to structural properties of mutualistic networks. (A) The temperature at which networks collapsed was plotted in relation to nestedness (NODF) for two levels of trait variation and two levels of heritability. Lines represent linear regression model with standard error. (B) Chances of abrupt collapse for all mutualistic networks in relation to nestedness for two levels of trait variation and two levels of trait heritability. Lines represent a generalised linear model with quasibinomial error distribution.

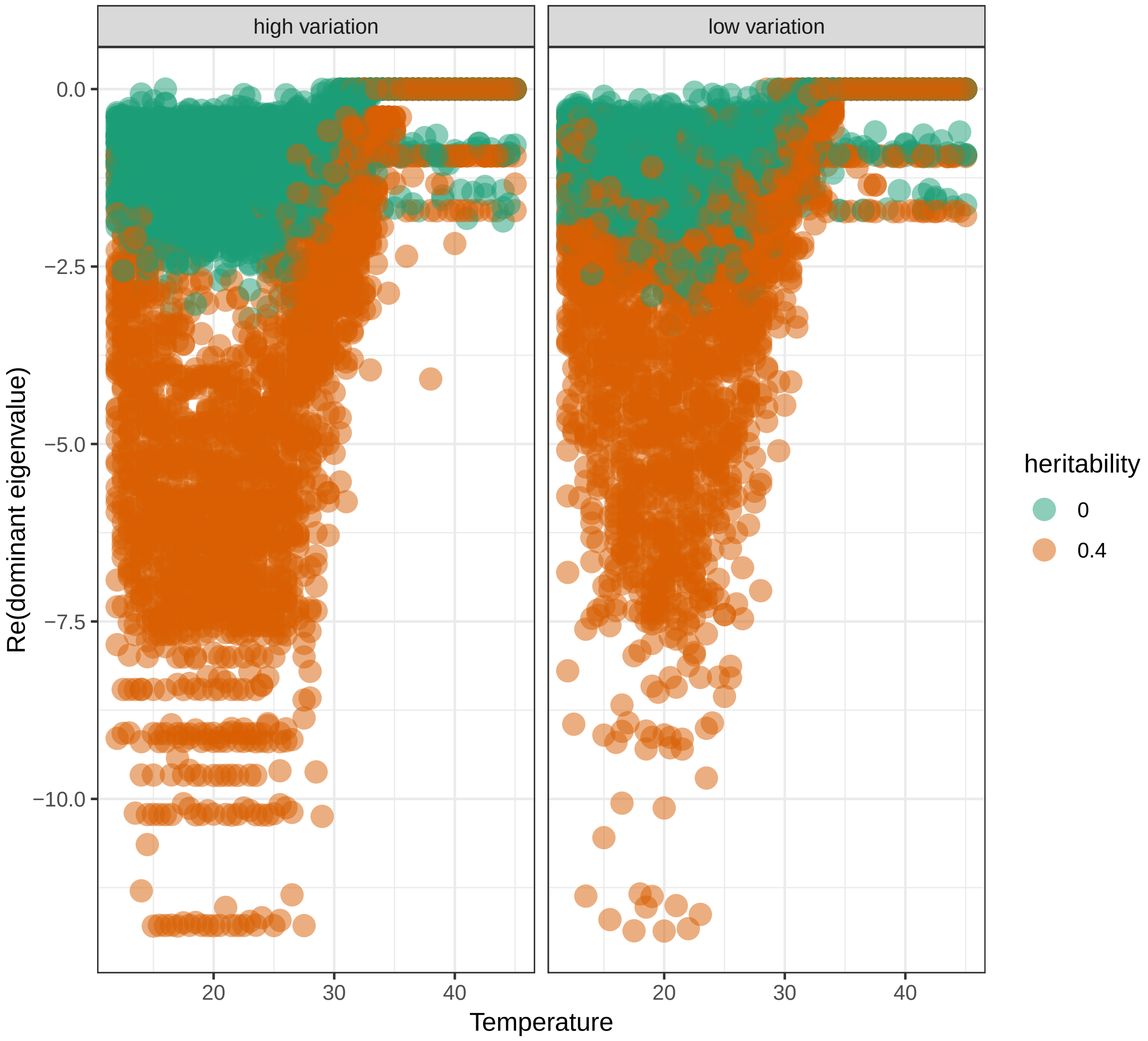


Figure S3: A) Local stability measured as real part of the dominant eigenvalue of the Jacobian matrix at equilibrium for 89 plant-pollinator networks in relation to local temperature for two levels of trait variation and heritability.

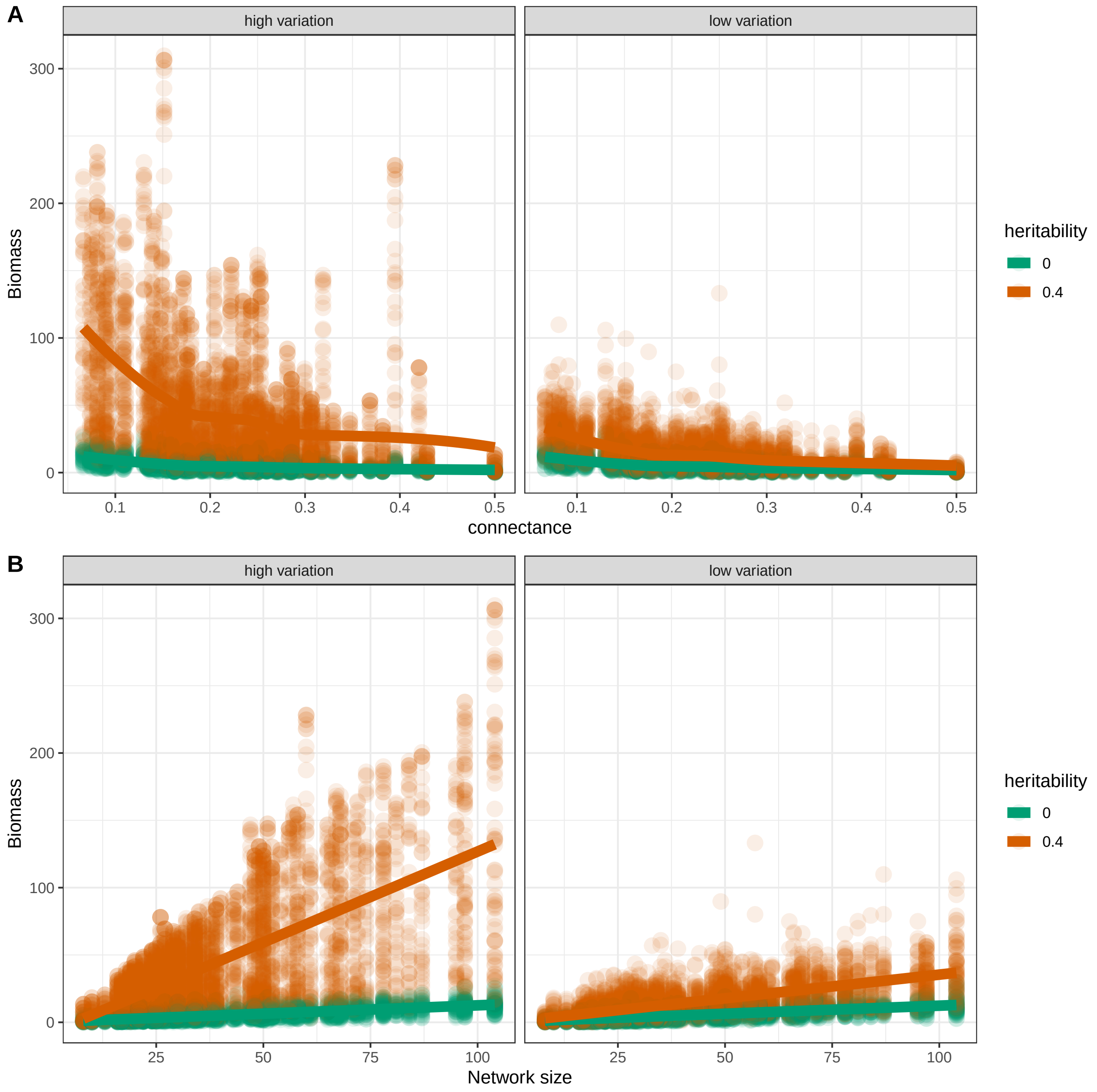


Figure S4: Equilibrium network biomass in relation to network connectance (A) and network size (B) grouped for local temperatures below $25^0C$.

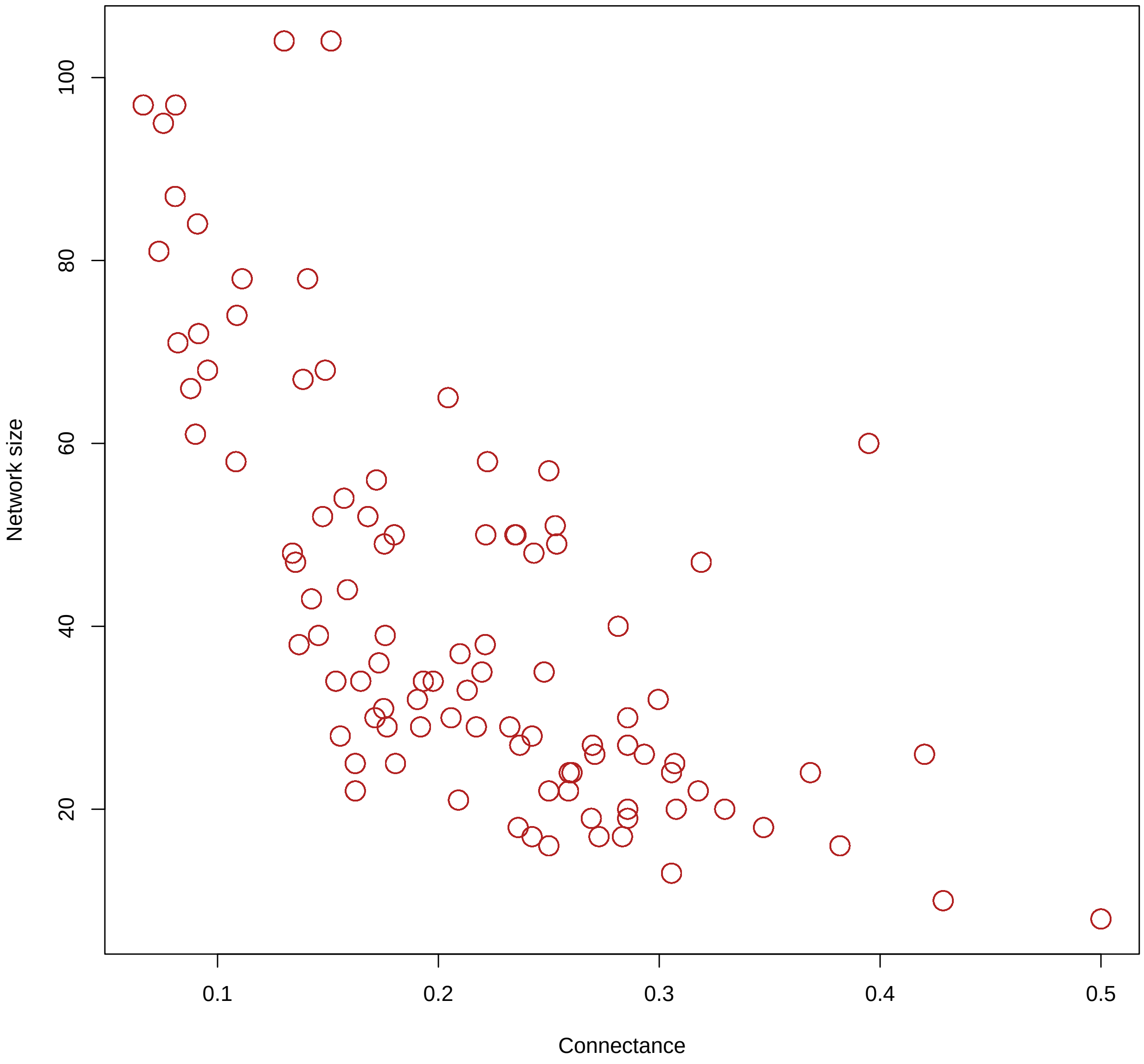


Figure S5: Relationship between network connectance and network size. Pearson correlation coefficient estimated to be -0.69.
